## Supplementary Figures for "BAZ2A-RNA mediated association with TOP2A and KDM1A represses genes implicated in prostate cancer"

### Suppl. Figure S1

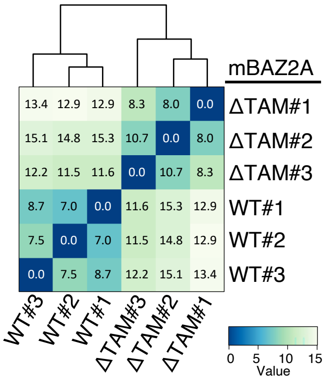


**Supplementary Figure S1**

Sample distances dendrogram of three BAZ2A$_{WT}$ samples (WT#1-3) and three BAZ2A$_{\Delta TAM}$ samples (ΔTAM #1-3) using Euclidean distance of the log$_2$-transformed counts from RNAseq. Dark colour indicates higher correlation between the samples.

### Suppl. Figure S2

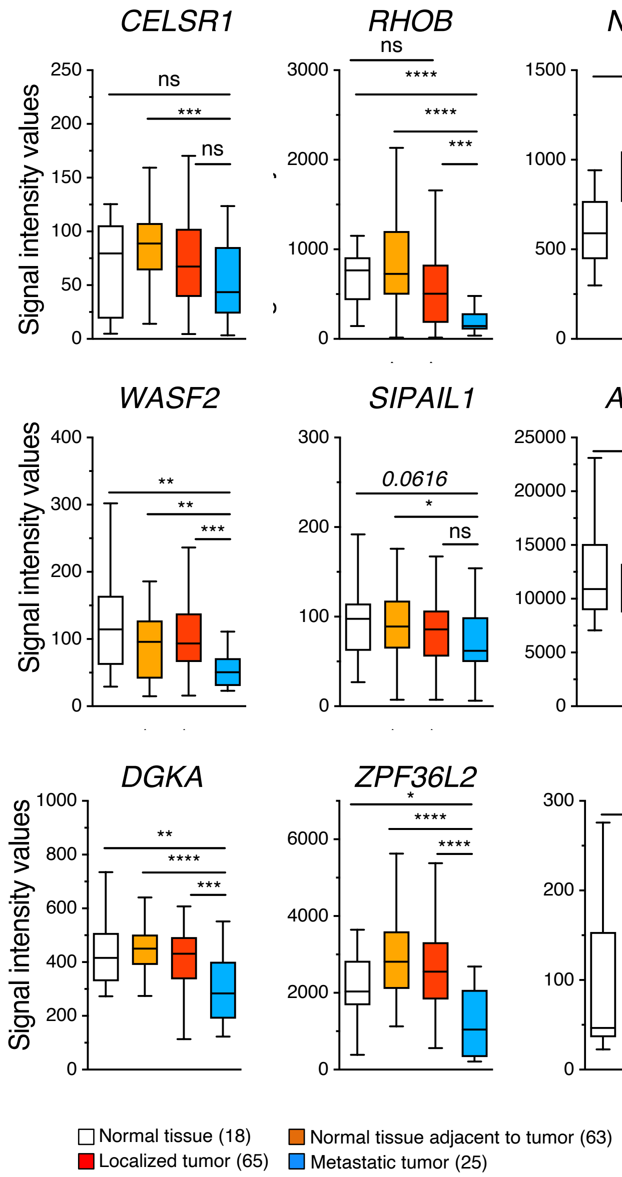


**Supplementary Figure S2**

Boxplots showing expression profiles of genes were linked to signal transduction, response to wounding and cell migration and motility that are repressed through BAZ2A-TAM domain from the gene expression microarray GEO data set GDS2545 (Chandran et al., 2007; Yu et al., 2004). Statistical significance (*P*-value) was calculated using two-tailed t-test (*<0.05, **<0.01, ***< 0.001, ****<0.0001); ns, not significant.

### Suppl. Figure S3

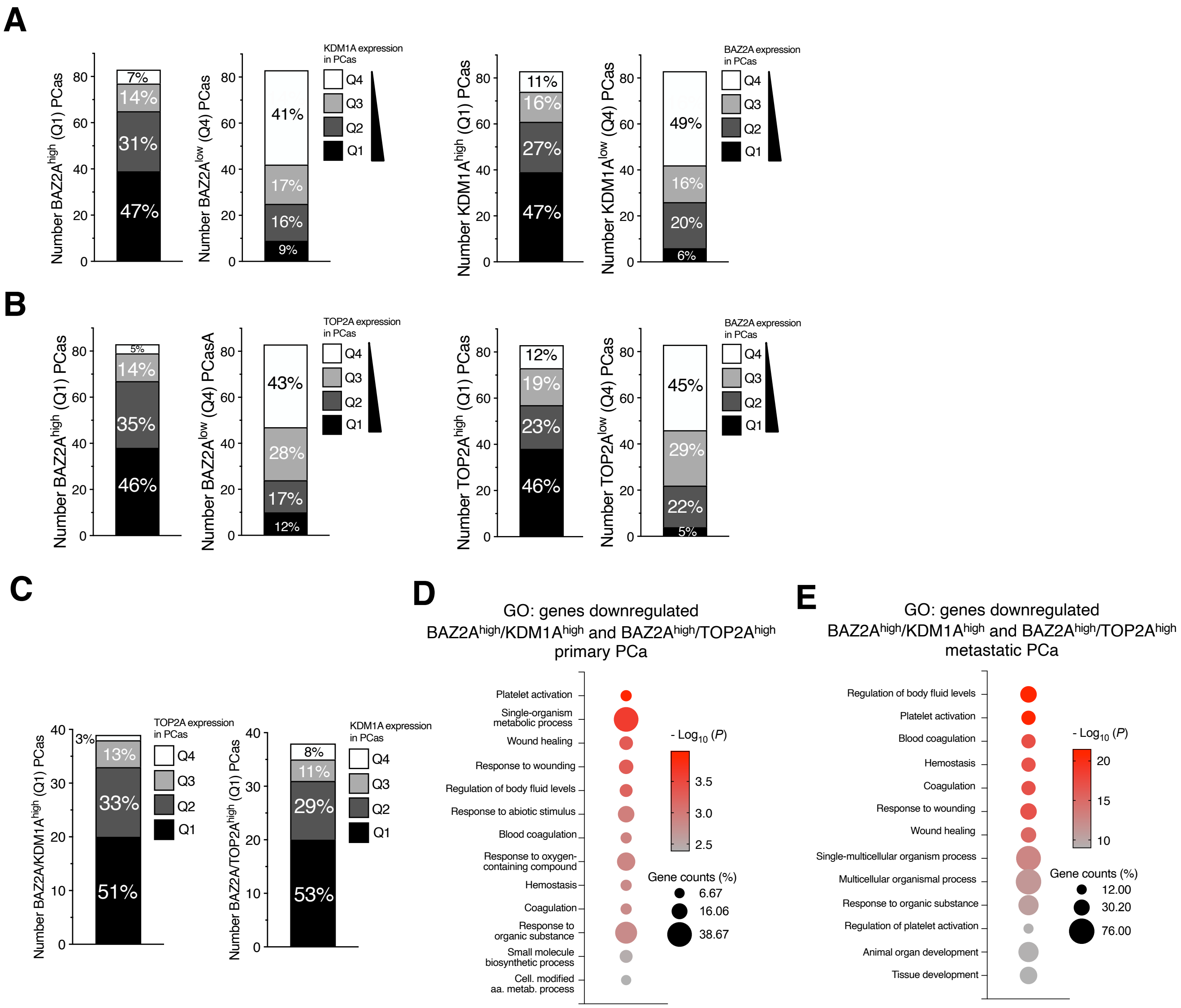


**Supplementary Figure S3**

**A.** Left panel. Quantifications of primary BAZ2A$^{high}$ and BAZ2A$^{low}$ PCas and the expression levels of KDM1A. Right panel. Quantifications of primary KDM1A$^{high}$ and KDM1A$^{low}$ PCas and the expression levels of BAZ2A. Quartile 1 (Q1): the top 25% of PCas with the highest expression. Quartile 4 (Q4): the top 25% of PCas with the lowest levels.

**B.** Left panel. Quantifications of primary BAZ2A$^{high}$ and BAZ2A$^{low}$ PCas and the expression levels of TOP2A. Right panel. Quantifications of primary TOP2A$^{high}$ and TOP2A$^{low}$ PCas and the expression levels of BAZ2A.

**C.** Left panel. Quantifications of primary BAZ2A$^{high}$/KDM1A$^{high}$and BAZ2A$^{low}$/KDM1A$^{low}$ PCas and the expression levels of TOP2A. Right panel. Quantifications of primary BAZ2A$^{high}$/TOP2A$^{high}$and BAZ2A$^{low}$/TOP2A$^{low}$ PCas and the expression levels of KDM1A.

**D.** Top gene ontology (GO) terms as determined using DAVID for genes downregulated in primary PCas BAZ2A$^{high}$/KDM1A$^{high}$ and BAZ2A$^{high}$/TOP2A$^{high}$.

**E.** Top gene ontology (GO) terms as determined using DAVID for genes downregulated in metastatic PCas BAZ2A$^{high}$/KDM1A$^{high}$ and BAZ2A$^{high}$/TOP2A$^{high}$.

### Suppl. Figure S4

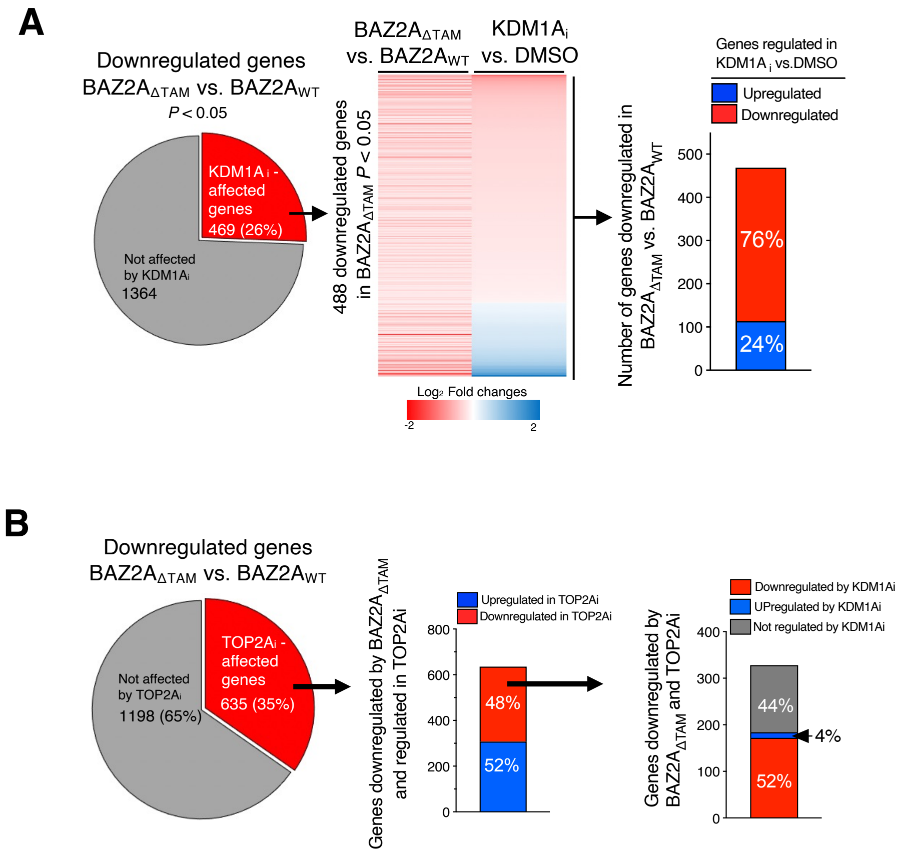


**Supplementary Figure 4**

**A.** Pie chart showing the number of genes significantly downregulated upon $BAZ2A_{\Delta TAM}$ expression in PC3 cells. In the middle panel is shown the heat map of $log_2$ fold change expression levels of genes downregulated by $BAZ2A_{\Delta TAM}$ and affected by KDM1Ai. In the right panel is shown the number of these genes that are significantly up- and downregulated by KDM1Ai.

**B.** Pie chart showing the number of genes significantly downregulated upon $BAZ2A_{\Delta TAM}$ expression in PC3 cells that are affected upon TOP2Ai treatment. Middle panel shows the number of genes downregulated by $BAZ2A_{\Delta TAM}$ and regulated by TOP2A that are up- and downregulated upon TOP2Ai treatment. The right panel shows the number of genes downregulated upon $BAZ2A_{\Delta TAM}$ expression and TOP2Ai treatment that are upregulated and downregulated upon KDM1Ai treatment.
